# Tetanus neurotoxin sensitive SNARE-mediated glial signaling limits motoneuronal excitability

**DOI:** 10.1101/2020.08.26.268011

**Authors:** Mathias A. Böhme, Anthony W. McCarthy, Monika Berezeckaja, Kristina Ponimaskin, Alexander M. Walter

**Author notes:** Correspondence to: Alexander M. Walter or Mathias A. Böhme. Editorial correspondence to, Dr. Alexander M. Walter, Molecular and Theoretical Neuroscience, Leibniz-Forschungsinstitut für Molekulare, Pharmakologie, Charité Campus Mitte, Charitéplatz 1, 10117 Berlin Germany.

## Abstract

Peripheral nerves contain motoneuron axons coated by glial cells, which essentially contribute to function but cellular reactions remain poorly understood. We here identify non-neuronal Synaptobrevin (Syb) as the essential vesicular SNARE in glia to insulate and metabolically supply *Drosophila* motoneurons. Interfering with Syb-functionality by glial knockdown, or glial expression of tetanus neurotoxin light chain (TeNT-LC) caused motonerve disintegration, blocked axonal transport, induced tetanic muscle hyperactivity and caused lethal paralysis. Surprisingly, not the established TeNT-LC-target, neuronal Synaptobrevin (nSyb), is the relevant SNARE, but non-neuronal Synaptobrevin (Syb): Knockdown of Syb- (but not nSyb-) phenocopied glial TeNT-LC expression whose effects were reverted by a TeNT-LC-insensitive Syb mutant. We link Syb-necessity to two distinct glia: to establish nerve insulating septate junctions in subperineurial glia and to integrate monocarboxylate transporters along the nerve in wrapping glia for motoneuronal metabolic supply. Our study identifies crucial roles of Syb in glial subtypes for nerve function and pathology, animal motility and survival.

## Introduction

Motor control is mediated by motoneurons of the central nervous system that can convey electrical information in the form of action potentials (APs) along their axons over large distances to peripheral body muscles. At the target muscle, motoneuronal APs induce the release of neurotransmitters (NT) at neuromuscular junctions (NMJs), leading to muscle excitation and contraction (Kuo and Ehrlich, 2015). Motoneuronal axons are bundled in nerves which also contain non-neuronal glial cells that separate motoneuron axons from the surrounding environment and additionally serve essential functions in development, regeneration, neural metabolism, ion homeostasis and AP propagation (Fields, 2015; Simard and Nedergaard, 2004; Zuchero and Barres, 2015). The functional importance of proper neural communication along peripheral nerves is evident from the many, often fatal diseases in which this is disrupted, including Charcot–Marie–Tooth disease, Guillain-Barré syndrome, amyotrophic lateral sclerosis, and tetanus (Bleck, 1989; Hardiman et al., 2017; Szigeti and Lupski, 2009; Willison et al., 2016).

The release of chemical transmitters, but also the delivery of proteins and lipids to different intracellular compartments depends on the fusion of cargo-containing vesicles with their target organelles. These fusion reactions are mediated by the formation of heat stable, coiled-coil soluble N-ethylmaleimide-sensitive-factor attachment receptor (SNARE) complexes between SNARE proteins resident on vesicle (v-SNARE) and target (t-SNARE) membranes (Bruns and Jahn, 2002; Jahn and Fasshauer, 2012; Jahn and Scheller, 2006). Synaptic transmission requires the fusion of neurotransmitter (NT) containing synaptic vesicles (SVs) with the plasma membrane which engages the evolutionarily conserved t-SNAREs syntaxin-1 and SNAP25 and the v-SNARE Synaptobrevin-2/VAMP2 (Syb2) in mammals or neuronal Synaptobrevin (nSyb) in *Drosophila* (DiAntonio et al., 1993; Jahn and Fasshauer, 2012). Tetanus neurotoxin (TeNT) is a potent bacterial toxin that abolishes NT release by TeNT-light chain (TeNT-LC) mediated cleavage of Syb-2/nSyb (Bruns et al., 1997; Schiavo et al., 1992; Schiavo et al., 2000; Sweeney et al., 1995) while the TeNT-heavy chain (TeNT-HC) targets the toxin to inhibitory interneurons of the vertebrate spinal cord (Blum et al., 2012; Blum et al., 2014; Bomba-Warczak et al., 2016; Deinhardt et al., 2006; Deinhardt and Schiavo, 2005; Erdmann et al., 1975; Lalli et al., 2003; Rummel, 2017; Schiavo et al., 2000; Schwab and Thoenen, 1976, 1978; Surana et al., 2018; Yeh et al., 2010). The resulting loss of inhibitory input onto spinal motoneurons causes hyperactivity, spastic paralysis and ultimately respiratory failure and death (Bleck, 1989; Popoff and Poulain, 2010; Surana et al., 2018). Despite the availability of functional vaccines, Tetanus still caused ~25000 death children in 2018 (Source: www.who.int) making a complete functional understanding of this disease essential. Interestingly, also glial cells, including the nerve-isolating peripheral Schwann cells, can take up TeNT (Huba and Hofmann, 1988; Schwab and Thoenen, 1978) and glial TeNT expression or disruption of its v-SNARE targets (VAMP2/Syb-2, VAMP3/cellubrevin) affects the function of central neurons (Lee et al., 2014; Pascual et al., 2005; Perea and Araque, 2007; Schwarz et al., 2017). However, whether TeNT in glial cells may affect motoneuroal function or even contribute to tetanus pathology remains unknown.

We here report on the unexpected observation that TeNT-LC expression in *Drosophila* glial cells severely disrupts peripheral nerve morphology and axonal transport of synaptic material. It furthermore caused motoneuronal hyperactivity and paralysis, typical effects of TeNT intoxication in higher organisms. Unlike neurons, where TeNT-LC cleaves nSyb and arrests neurotransmission, in glial cells these effects were caused by the functional loss of non-neuronal Syb and could be rescued by a TeNT-insensitive Syb-variant. TeNT-LC expression in axon encircling wrapping glia (WG) reduced monocarboxylate transporters at the glia/neuronal interface and caused an axonal accumulation of the synaptic protein Bruchpilot (BRP). Similar aberrant axonal transport was observed upon WG knockdown of Basigin, a protein that targets monocarboxylate transporters to plasma membranes, suggesting that these effects are due to a disruption in glia-to-neural metabolic supply. In contrast, TeNT-LC (or Syb-RNAi) expression in subperineurial glial cells additionally disrupted nerve morphology as well as septate junction formation and in some cases caused aberrant nerve activity. In conclusion we report on SNARE-mediated reactions in glial cells that are essential for motor function and whose disruption may cause severe pathology, including tetanus.

## Results and discussion

### Expression of TeNT-LC in glial cells disrupts nerve morphology and function

Classic experiments revealed TeNT uptake into Schwann cells (Schwab and Thoenen, 1978) and glial cells exocytose substances influencing neural function possibly in a SNARE-dependent manner (Carlsen and Perrier, 2014; Christensen et al., 2018; Christensen et al., 2013; Gucek et al., 2012; Schwarz et al., 2017; Verderio et al., 2012). We therefore wondered whether TeNT-LC targeted to glial cells could also affect motoneuronal function.

Three types of *Drosophila* glial cells are found in a compound organization coating the segmentally arranged abdominal peripheral nerves (Bittern et al., 2020; Stork et al., 2008) and Fig. 1A,B), which harbor the sensory- and motoneuron axons (motoneuronal innervation depicted in Fig 1B, left). We started our investigation by expressing TeNT-LC (Sweeney et al., 1995) in all glial subtypes using the pan-glial driver Repo-Gal4 (Sepp et al., 2001) and investigated the influence on larval behavior, peripheral nerve morphology (via staining with horse-radish peroxidase (HRP) as neuronal membrane marker) and function (Fig. 1C-H). Surprisingly, pan-glial TeNT-LC expressing larvae were severely paralyzed, hardly able to move and died at late larval stages (Fig 1C,D and Video 1-3). Additionally, peripheral nerves of pan-glial TeNT-LC expressing larvae were morphologically severely disrupted and showed partial defasciculation (Fig. 1E, arrow head for single axon leaving the nerve trunk) and an expansion of the nerve area (Fig. 1E,F). A recent study reported similar, but locally constrained nerve swellings upon glial knockdown of the salt-inducible kinase 3 (SIK3), a central node in a signal transduction pathway controlling glial K^+^ and water homeostasis (Li et al., 2019). In contrast, peripheral nerves of pan-glial TeNT-LC expressing animals showed a continuous disruption of the entire axonal length although to different degrees. We quantitatively evaluated the severity of the disruption by categorizing the nerves by their degree of disintegration (<u>normal</u>: long, thin and smooth nerves (see control images in Fig. 1E for example); <u>intermediate</u>: slight morphological alterations like single defasciculating axons, slightly increased nerve area; <u>disrupted</u>: complete disintegration of the nerve, defasciculation of whole parts of the nerve, large expansion of the nerve area). While control groups (larvae with either the Repo-Gal4 driver or the UAS-TeNT-LC construct alone) almost exclusively showed normal nerve morphologies, most nerves expressing pan-glial TeNT-LC were disrupted (Fig. 1F, nerve morphology). Additionally, pan-glial TeNT-LC expression caused a five-fold increase in the number of spots in the axon containing presynaptic protein BRP (labelled by immunostaining; Fig 1E,F), indicating a disruption in the synaptic delivery of this protein.

**Figure 1:**
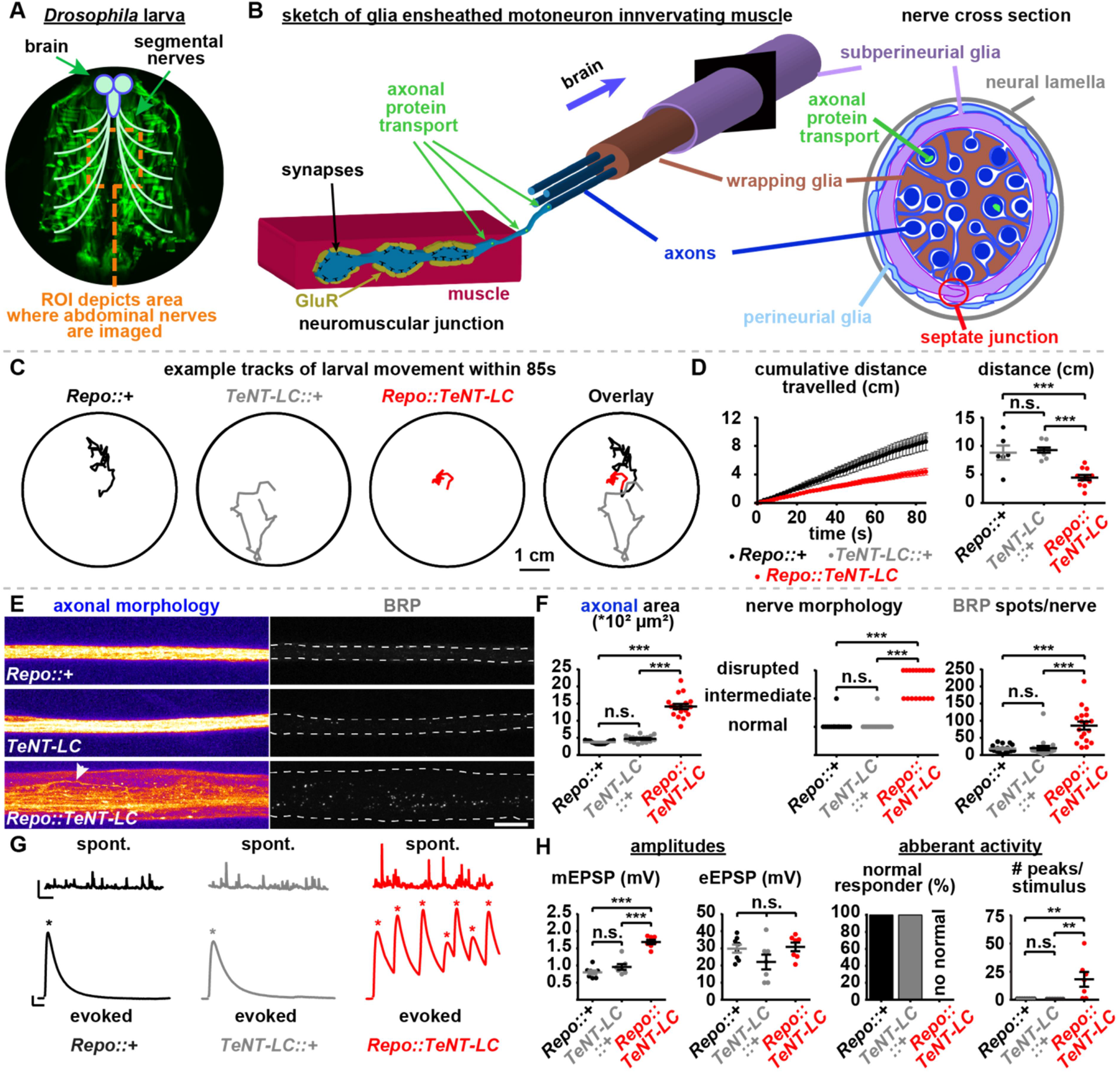
Pan-glial TeNT-LC expression causes paralysis, disrupts nerve integrity, and results in motoneural hyperactivity. **A,B** Sketches of dissected larvae depicting segmental nerves radiating from the brain (A) containing muscle innervating, glia ensheathed motoneurons (B, left; nerve cross section, right, according to (Stork et al., 2008)). **C,D** Analysis of larval movement with example tracks over 85s (C), distances travelled over time (85s; D, left) and comparison of final distances travelled by 3^rd^ instar larvae of the indicated genotypes (D, right): *Repo::*+ (black), *TeNT-LC::*+ (grey) and *Repo::TeNT-LC* (red). See also Videos 1-3. **E,F** Nerves of segments A2–A4 (E) and quantification of axonal area, nerve morphology and BRP spots per nerve (F) were investigated in the region of interest (ROI) illustrated in panel A in 3^rd^-instar larvae of *Repo::*+ (black), *TeNT-LC::*+ (grey) and *Repo::TeNT-LC* (red) animals. Dashed line in (E, BRP) indicates nerve area and arrow head indicates defasciculated single axon. **G,H** Representative mEPSP (spont.) and eEPSP (evoked) traces (G) and quantification of mEPSP/eEPSP amplitudes, % of recorded cells displaying normal activity (single eEPSP in response to single stimulation) and the number of response peaks per stimuli in *Repo::* + (black), *TeNT-LC::*+ (grey) and *Repo::TeNT-LC* (red) animals. Asterisks indicate postsynaptic response peaks (eEPSP). Scale bars: (C) 1 cm; (E) 10 μm; (G) mEPSP: 1 s, 2 mV. eEPSP: 25 ms, 5 mV. Statistics: parametric one-way analysis of variance (ANOVA) test, followed by Tukey’s multiple comparison test except for (F, nerve morphology) where a non-parametric Kruskal-Wallis test was performed. ***p ≤ 0.001; **p ≤ 0.01; *p ≤ 0.05; n.s. (not significant) p > 0.05. (C,D) *Repo::*+: six larvae; *TeNT::*+: nine larvae; *Repo::TeNT:* 11 larvae; (E,F) *Repo::*+: 17 nerves, six larvae; *TeNT::*+: 18 nerves, six larvae; *Repo::TeNT:* 18 nerves, six larvae; (G,H) *Repo::*+: 8 cells, four larvae; *TeNT::*+: 7 cells, four larvae; *Repo::TeNT:* 8 cells, four larvae. All panels show mean ± s.e.m.

Peripheral nerves contact body wall muscles at NMJs (Fig. 1B) where NTs are released by motoneuron upon APs which induce depolarization of the muscle membrane potential, contraction, and finally movement. To investigate whether glial expression of TeNT-LC affected this we performed recordings of muscle’s membrane potential (Fig. 1G,H). Without stimuli, spontaneous NT release gives rise to ‘miniature’ excitatory postsynaptic potentials (mEPSPs) and the amplitudes of these mEPSPs were almost doubled upon pan-glial TeNT-LC expression compared to controls (Fig. 1G,H, spont.) while their frequency was similar (*Repo*:: +: 2.054 ± 0.1678 Hz, *TeNT-LC*:: +: 2.538 ± 0.2536 Hz, *Repo::TeNT-LC*: 1.874 ± 0.1812 Hz; one-way ANOVA with Tukey’s multiple comparison test, *Repo*:: + vs. *TeNT-LC*::+: p = 0.226, *Repo*::+ vs. *Repo*::*TeNT-LC*: p = 0.8009, *TeNT-LC*::+ vs. *Repo*::*TeNT-LC*: p = 0.0837). Electrical stimulation of the innervating nerve in control animals evokes a single AP that reliably triggered a single evoked excitatory postsynaptic potential in the muscle (eEPSP; seen in 8/8 driver control and 7/7 TeNT-LC-construct control cells; Fig. 1G,H, evoked). In contrast, upon pan-glial TeNT-LC expression, a single stimulus triggered additional eEPSPs and thus motoneuronal hyperactivity (ranging from 2 to 246 additional events per cell, seen in 10/10 cells; Fig. 1G,H), a typical hallmark of TeNT intoxication in higher organisms. This is a similar observation to one seen with glial-knockdown of SIK3, although in that case supernumerary EPSPs also occurred without any stimulation (Li et al., 2019). While the number of eEPSPs elicited per stimulus was increased, the average amplitude of the first eEPSPs we observed did not differ from control cells (Fig. 1G,H), indicating that glial TeNT expression did not disrupt synaptic transmission *per se*. Thus, we report that TeNT-LC-mediated interference with SNARE-dependent processes in glial cells leads to paralysis, disrupts nerve integrity, impairs axonal transport, and causes motoneural hyperexcitability.

### Essential glial functions are mediated by Syb but not nSyb

We next asked which v-SNARE might be targeted by TeNT-LC in glial cells (Fig. 2A). Tetanus pathology is caused by Synaptobrevin-2 cleavage in mammalian interneurons (Schiavo et al., 2000). However, TeNT-LC also cleaves other v-SNAREs, including VAMP1/Synaptobrevin-1 enriched in spinal cord neurons and the more ubiquitously expressed VAMP3/cellubrevin in mammals (Carle et al., 2017; Elferink et al., 1989; McMahon et al., 1993; Patarnello et al., 1993; Schiavo et al., 1992). The *Drosophila* genome harbors two putative TeNT-LC targets, neuronal-Synaptobrevin (nSyb) which mediates SV fusion (Deitcher et al., 1998; DiAntonio et al., 1993; Sweeney et al., 1995) and the rather ubiquitously expressed synaptobrevin (Syb) which is likely involved in more constitutive vesicular fusion reactions (Chin et al., 1993; Sudhof et al., 1989). TeNT-LC is thought to specifically cleave *Drosophila* nSyb (Sweeney et al., 1995). However, an alignment of nSyb and non-neuronal Syb revealed differences mainly in their N- and C-terminal parts while the central SNARE-motif (Fig. 2B, yellow shaded) is largely conserved (Fig. 2B). Importantly, the glutamine (Q) – phenylalanine (F) TeNT-LC-cleavage site (Schiavo et al., 1992), is conserved in both proteins (Fig. 2B, red box).

**Figure 2:**
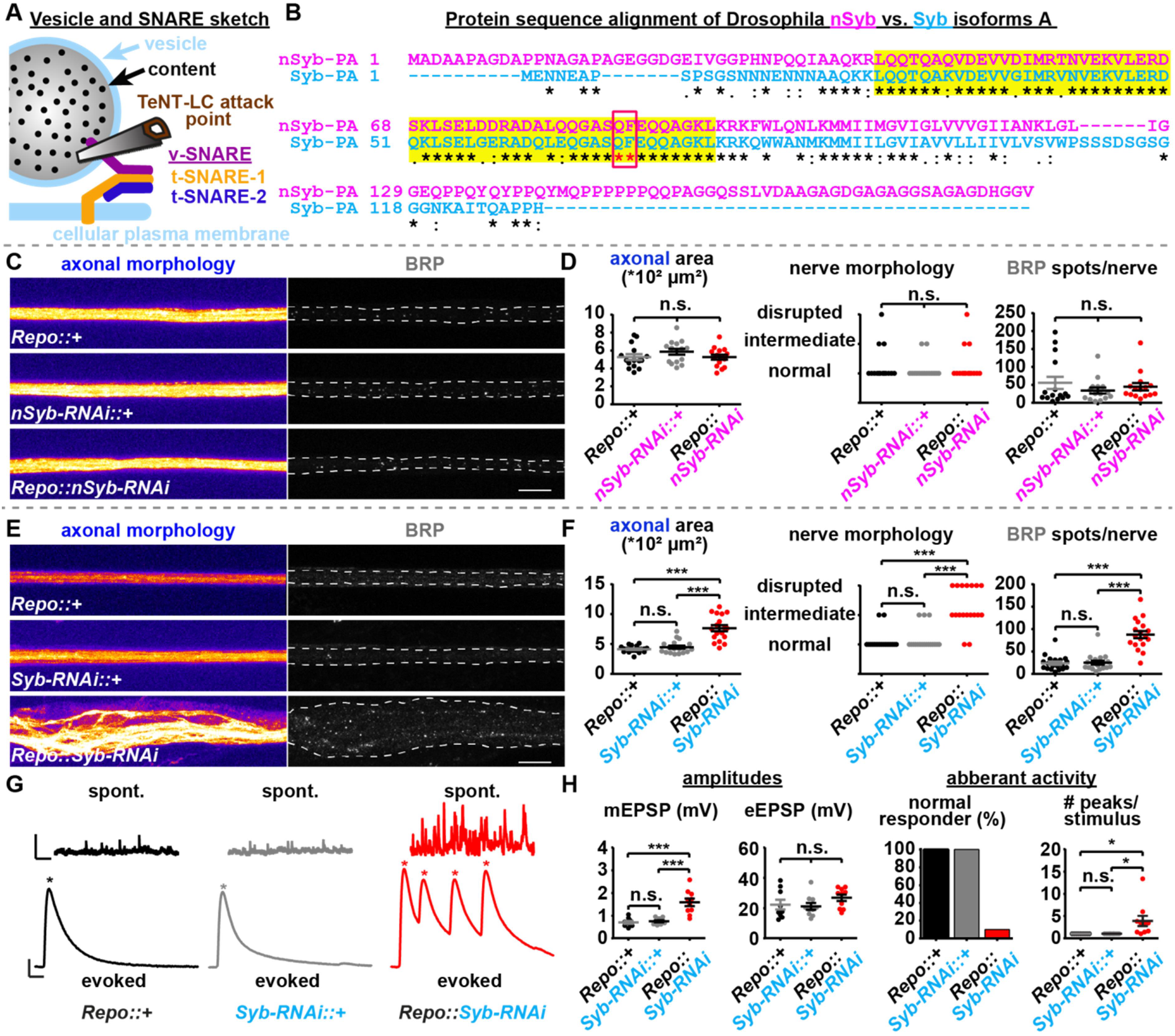
Syb but not nSyb is targeted by TeNT-LC in glial cells. **A** Sketch of a vesicle depicting the SNARE proteins and the v-SNARE attack point of TeNT-LC. **B** Sequence alignment of Isoforms A of *Drosophila* nSyb (magenta) and Syb (blue). SNARE motif (yellow shaded) and the TeNT-LC cleavage site (QF, red box) are highlighted. Alignment performed using Clustal Omega. Gonnet PAM250 matrix used to compare sequence substitutions: * identical aa,: score >0.5,. score <0.5, “gap” score below 0. **C-F** Nerves of segments A2–A4 (C,E) and quantification of axonal area, nerve morphology and BRP spots per nerve (D,F) from 3^rd^-instar larvae of the indicated genotypes. Dashed line in C,E indicates nerve area. **G,H** Representative mEPSP (spont.) and eEPSP (evoked) traces (G) and quantification of mEPSP/eEPSP amplitudes, % of recorded cells displaying normal activity (single eEPSP in response to single stimulation) and the normalized number of response peaks per stimulus in *Repo::+* (black), *Syb-RNAi:: +* (grey) and *Repo::Syb-RNAi* (red) animals. Asterisks indicate postsynaptic response peaks (eEPSP). Scale bars: (C, E) 10 μm; (G) mEPSP: 1 s, 2 mV. eEPSP: 25 ms, 5 mV. Statistics: parametric one-way analysis of variance (ANOVA) test, followed by Tukey’s multiple comparison test except for (D, F: nerve morphology) where a non-parametric Kruskal-Wallis test was performed. ***p ≤ 0.001; *p ≤ 0.05; n.s. (not significant) p > 0.05. (C,D) *Repo::*+: 15 nerves, six larvae; *nSyb-RNAi::*+: 15 nerves, six larvae; *Repo::nSyb-RNAi:* 15 nerves, six larvae; (E,F) *Repo::* +: 19 nerves, six larvae; *Syb-RNAi::*+: 18 nerves, six larvae; *Repo::Syb-RNAi:* 19 nerves, six larvae; (G,H) *Repo::*+: 9 cells, five larvae; *Syb-RNAi::*+: 10 cells, five larvae; *Repo::Syb-RNAi: 10* cells, five larvae. All panels show mean ± s.e.m.

We thus sought to investigate whether disruption of Syb or nSyb was causative for the observed defects upon TeNT-LC expression by testing whether pan glial (Repo-Gal4) knockdown of either protein using specific RNAi-lines induced similar effects (Fig. 2C-H). Pan-glial nSyb knockdown did not change nerve morphology, nerve area or on axonal BRP presence in comparison to controls (Fig. 2C,D; similar results were obtained with a second nSyb-RNAi line (#VDRC 49201, data not shown)). In contrast, pan-glial knockdown of non-neuronal Syb phenocopied pan-glial TeNT-LC expression with largely increased nerve areas, severely disrupted nerve morphologies and enhanced axonal BRP accumulations (Fig. 2E,F). Similarly to glial TeNT-LC expression, mEPSP amplitudes were increased and cells exhibited hyperactive motoneuronal behavior with supernumerary eEPSPs after stimulation (ranging from 2 to 62 additional events per cell, seen in 9/10 cells; Fig. 2G,H) but similar eEPSP amplitudes (Fig. 2G,H). Thus, TeNT-LC-mediated impairment of Syb- and not nSyb-function in *Drosophila* glial cells is likely responsible for disrupted nerve integrity, axonal accumulation of synaptic proteins and hyperactive motoneuronal responses (see also below).

### Syb disruption in subperineurial glia causes paralysis, distorts nerves and impairs axonal transport

We next speculated how TeNT-LC or Syb-knockdown cause these phenotypes and whether all glial subtypes are affected. *Drosophila* peripheral axons are supported by three glial types: wrapping glia (WGs), subperineurial glia (SPGs) and perineurial glia (PGs) ((Stork et al., 2008) and Fig. 1B). SPG enwrap axons and WG ((Stork et al., 2008) and Figs. 1B & 3A) and additionally extend to fully cover the NMJ in 35% of cases (investigated using rl82-Gal4 (Gli-Gal4 (Auld et al., 1995; Sepp and Auld, 1999)) mediated expression of membrane associated GFP (mCD8-GFP); Fig. 3B,C), in line with previous results (Auld et al., 1995; Brink et al., 2009; Fuentes-Medel et al., 2009; Kerr et al., 2014; Sepp and Auld, 1999).

**Figure 3:**
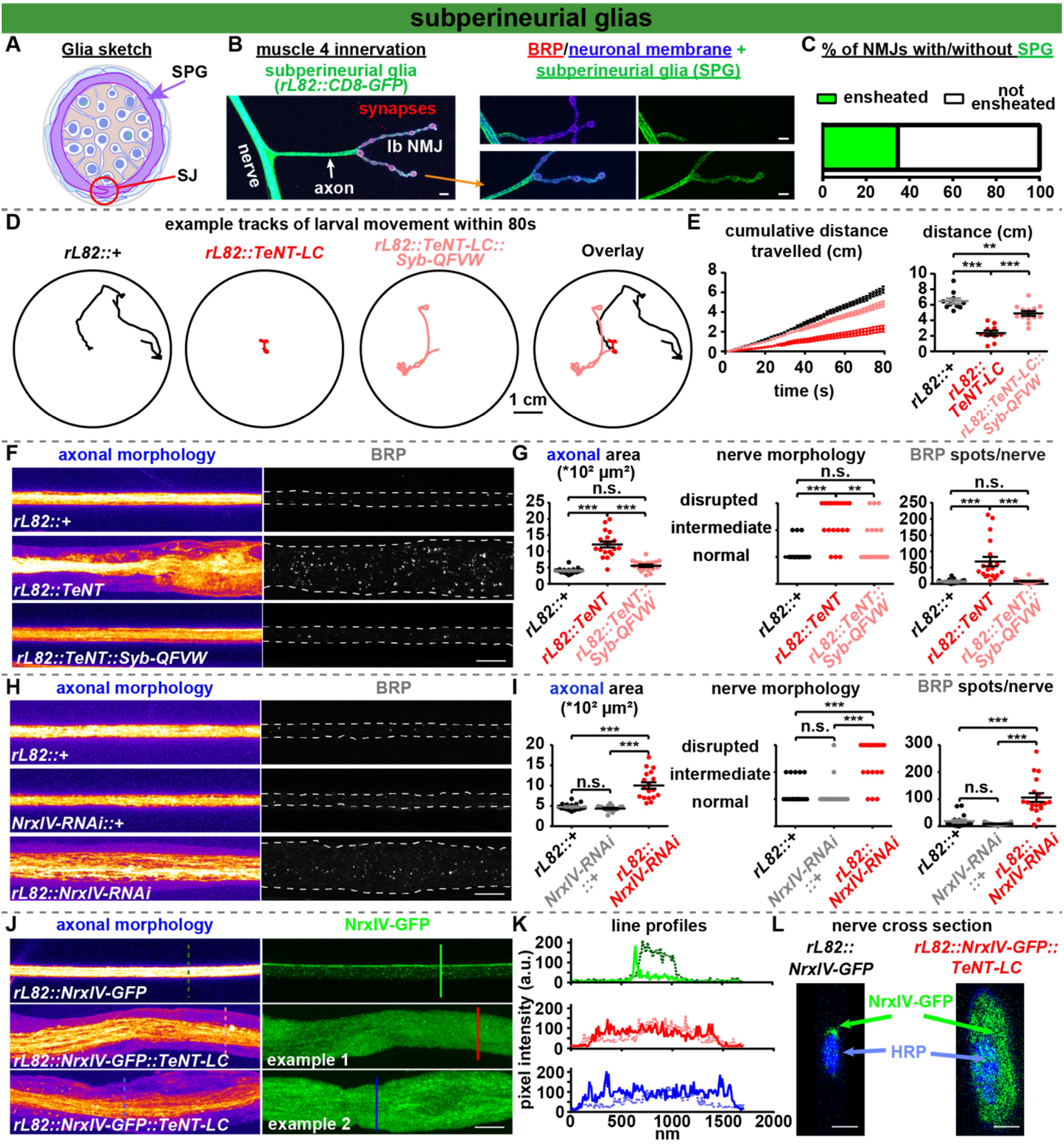
TeNT-LC targets Syb in SPGs and impairs locomotion, disrupts peripheral nerves and axonal transport, and blocks SJ formation. **A** Cross section sketch of glial ensheathed motoneuron axon highlighting SPGs (purple). Red circle highlights septate junction (SJ). **B,C** Examples of muscle 4 NMJs of segments A2–A4 from 3^rd^-instar larvae expressing CD8-GFP in SPGs (left) with (left & bottom) and without (right top) SPG ensheathment of the NMJ. **C** Quantification of % of NMJs with/without SPG ensheatment. **D,E** Analysis of larval movement with example tracks over 80s (D), distances travelled over time (80s; E, left) and comparison of final distances travelled by larvae of the indicated genotypes (E, right). See also Videos 4, 5 and 7. **F,G** Nerves of segments A2–A4 (F) and quantification of axonal area, nerve morphology and BRP spots per nerve (G) from 3^rd^-instar larvae of *rL82::*+ (black), *rL82::TeNT* (red) and *rL82::TeNT::Syb-QFVW*(light red) animals. **H,I** Same as in F-G but for *rL82::*+ (black), *NrxIV-RNAi::*+ (grey) and *rL82::NrxIV-RNAi* (red) animals. **J-L** Nerves of segments A2–A4 (J), vertical line profiles across nerve NrxIV-GFP signals (K; solid line GFP signal, dashed line HRP signal) and orthogonal nerve cross section (L) from 3^rd^-instar larvae of *rL82:: NrxIV-GFP* and *rL82::NrxIV-GFP::TeNT* (two examples are shown). Lines in J indicate line profile positions shown in K. Scale bars: (B) 5 μm; (F,H,J) 10 μm; (L) 2.5 μm. Statistics: parametric one-way analysis of variance (ANOVA) test, followed by Tukey’s multiple comparison test except for (G, I: nerve morphology) where a non-parametric Kruskal-Wallis test was performed. ***p ≤ 0.001; **p ≤ 0.01; *p ≤ 0.05; n.s. (not significant) p > 0.05. Dashed lines in (F,H: BRP) indicate nerve area. (B,C) 26 NMJs, seven larvae; (D,E) *rl82::*+: ten larvae; *rL82::TeNT:* ten larvae; *rL82:TeNT::Syb-QFVW:* 13 larvae; (F,G) *rl82::* +: 17 nerves, six larvae; *rL82::TeNT:* 19 nerves, five larvae; *rL82:TeNT::Syb-IsoA-QFVW:* 18 nerves, six larvae; (H,I) *rL82::+:* 17 nerves, six larvae; *NrxIV-RNAi::*+: 17 nerves, six larvae; *rL82::NrxIV-RNAi:* 18 nerves six larvae; (J-L) *rL82::NrxIV-GFP:* 18 nerves, six larvae; *rL82::NrxIV-GFP::TeNT:* 18 nerves, six larvae. All panels show mean ± s.e.m. See also Figs. S1 and 2.

Similarly to pan-glial TeNT-LC expression (Fig. 1C,D and Videos 1-3) SPG TeNT-LC expressing larvae showed severely reduced locomotion (Fig. 3D,E and Videos 4,5), displayed distorted nerve morphology and axon BRP accumulation (Fig. 3F,G). In the course of this study we also found two rare adult escapees of TeNT-LC expressing SPGs which were also severely paralyzed, showed almost no movement and were even unable to re-orientate to an upright stance when falling on their back (Video 6). To verify that Syb was the relevant TeNT target, we generated a TeNT-insensitive Syb variant (as UAS-construct of Syb Isoform A) by exchanging the glutamine (Q_69_) – phenylalanine (F_70_) TeNT-LC-cleavage site (Fig. 2B) with Valin (V) – Tryptophan (W) (Regazzi et al., 1996). Co-expression of this TeNT-insensitive UAS-Syb-QFVW in SPGs almost completely restored locomotion, and rescued nerve morphology and axonal BRP accumulation (Fig. 3D-G and Video 7). Additionally, SPG Syb-knockdown also elicited nerve disintegration and axonal BRP accumulation, clearly implicating Syb as the relevant TeNT target in glial cells (Fig. S1A-C).

Electrophysiological recordings of larvae expressing both SPG TeNT-LC or Syb-RNAi revealed occasional aberrant motoneuronal responses after AP-evoked stimulation (Fig. S1D-G) but these effects were much weaker than upon pan-glial expression which might be due to the additional contribution of other glial cell types (e.g. wrapping glia, (Li et al., 2019)) or due to a weaker expression strength of the rL82-Gal4 driver compared to Repo-Gal4 (Yildirim et al., 2019). We can exclude that TeNT-LC or Syb-RNAi expression kills SPGs cells as their visualization by simultaneous CD8-GFP expression showed an intact GFP coverage of investigated nerves similar to control animals (Fig. S2A-D).

We next wondered what biological processes may be disrupted upon TeNT-LC expression in SPGs. By coincidence we discovered that an antibody staining the glutamate receptor subunit IID (GluRIID, (Qin et al., 2005)) reliably labelled an outer layer of the nerve, possibly the neural lamella. Though the epitope responsible is unknown, GluRIID staining revealed extensive lamellar folds and large “voids” within the axonal HRP staining in larvae expressing Syb-RNAi in SPGs (Fig. S1A,B; neural lamella). These “voids” together with the morphological nerve phenotypes (disintegration, enlarged nerve area) are qualitatively similar to observations in mutants that interfere with glial wrapping of peripheral axons or with the formation of septate junctions (SJs) (Babatz et al., 2018; Leiserson et al., 2000). These SJs seal off the axon/WG encircling SPGs (Fig 1B, right) to build an occluding barrier for metabolic insulation (Banerjee et al., 2008; Baumgartner et al., 1996; Carlson et al., 2000; Schwabe et al., 2005; Stork et al., 2008; Yildirim et al., 2019). We therefore analyzed how disruption of SJ formation affected peripheral nerve morphology by interfering with a critical SJ component, the transcellular adhesion protein Neurexin IV (NrxIV; (Babatz et al., 2018; Baumgartner et al., 1996; Oshima and Fehon, 2011; Stork et al., 2008)). Remarkably, very similar to TeNT/Syb-RNAi expression, NrxIV-RNAi knockdown in SPGs disrupted nerves, increased their area and caused axonal BRP-accumulation (6-fold increase in comparison to controls; compare Fig. 3H,I with 3F,G and Fig. S1A-C). Thus, interference with Syb or SJ formation in SPGs causes similar effects.

SJ formation depends on the delivery of key components (including NrxIV) to the glial surface likely by exocytosis (Babatz et al., 2018; Tiklova et al., 2010) and we hypothesized that this was driven by Syb. We therefore tested whether TeNT-LC expression interfered with NrxIV delivery to the SJ by studying its cellular localization using an endogenous GFP-tag (NrxIV-GFP; (Edenfeld et al., 2006)). In control nerves, NrxIV-GFP was very restricted to a linear profile along the peripheral nerves (Fig. 3J-L; note peak in vertical nerve line profil (Fig. 3J,K; top panel, solid green line) and orthogonal nerve corss-sections revealed a single large GFP signal, indicating proper SJ formation (Fig. 1B; 3L, left). Remarkably, TeNT expression led to a diffuse, rather unspecific NrxIV-GFP distribution througout the nerve, suggesting a loss of the SJ (Fig. 3J,K bottom panels for two examples and 3L, right). Our resutls are consistent with an essential function of Syb in SJ formation which partially explains the deteriotations observed upon Syb interference.

Electrophysiological characterization of NrxIV-RNAi expressing SPGs revealed increased mEPSP and eEPSP amplitudes (Fig. S2E,F) but unlike pan-glial TeNT-LC/Syb RNAi expression no hyperactivity was seen (Fig. S2E,F). This could again reflect the involvement of additional glial subtypes or a less severe disruption of SJ formation than upon pan-glial TeNT-LC/Syb-RNAi expression. Notably, the impaired blood-brain barrier function in *nrxIV* mutants is partially compensated by the formation of intertwined cell-cell protrusions, resembling an evolutionary ancient barrier type found in primitive vertebrates or invertebrates (Babatz et al., 2018; Bundgaard and Abbott, 1992, 2008; Stork et al., 2008) potentially sufficient to suppress hyperexcitability. Alternatively or additionally, TeNT-LC/Syb-RNAi expression might impair other exocytotic processes (Hoogstraaten et al., 2020; Pascual et al., 2005; Perea and Araque, 2007; Schwarz et al., 2017). Recently, interference with WNT- and thus peptidergic-signaling specifically in SPGs was shown to affect postsynaptic glutamate receptors and synaptic transmission (Kerr et al., 2014). Indeed, Syb-RNAi expression in SPGs caused an accumulation of the peptidergic vesicle marker atrial natriuretic peptide (ANF (Rao et al., 2001); 11-fold increase in comparison to control; Fig. S2G,H) suggesting that peptidergic release from SPG is also mediated by Syb and might contribute to nerve function. In conclusion, the disruption of nerves upon TeNT-LC expression in SPGs appears to be due to a block of Syb-mediated SJ formation. However, the prominent motoneural hyperactivity seen upon pan-glial expression is likely due to the disruption of additional secretory processes, possibly in other cell types.

### TeNT-LC expression in WG impairs axonal transport likely by disrupting neural metabolic supply

SPG cells are not in direct contact with peripheral axons after early larval stages (Fig. 1B and (Stork et al., 2008)). We thus wondered whether the influence of SPGs on axonal transport might be indirect and rather via WGs whose functionality might also be impaired when nerve integration or signaling from the SPGs is disrupted. Although WGs directly contact axons/axon bundles, they rarely extend to the NMJ and we did not observed them to fully cover it (Fig. 4A-C). TeNT-LC expression in WGs using the WG-specific Nrv2-Gal4 driver line (although expression of Nrv2-Gal4 in cortex glia was also reported, which, however, are not present at peripheral nerves (Stork et al., 2008; Yildirim et al., 2019)) did not kill WG cells examined via simultaneous CD8-GFP expression (Fig. S3A,B). Unlike TeNT-LC expression in SPGs, TeNT-LC expression in WGs neither disrupted nerve integrity nor altered nerve area but caused a similarly strong accumulation of axonal BRP (Fig. 4D,E). Thus, axonal accumulation of synaptic BRP can be genetically uncoupled from the nerve defasciculation and hyperexcitablity described above. Accordingly, the none-additive nature of the effects could imply that axonal BRP accumulations observed upon SPG-perturbation (Figs. 3F,G,H,I; Fig. S1A-C) may be caused by an indirect interference with WG function.

**Fig. 4:**
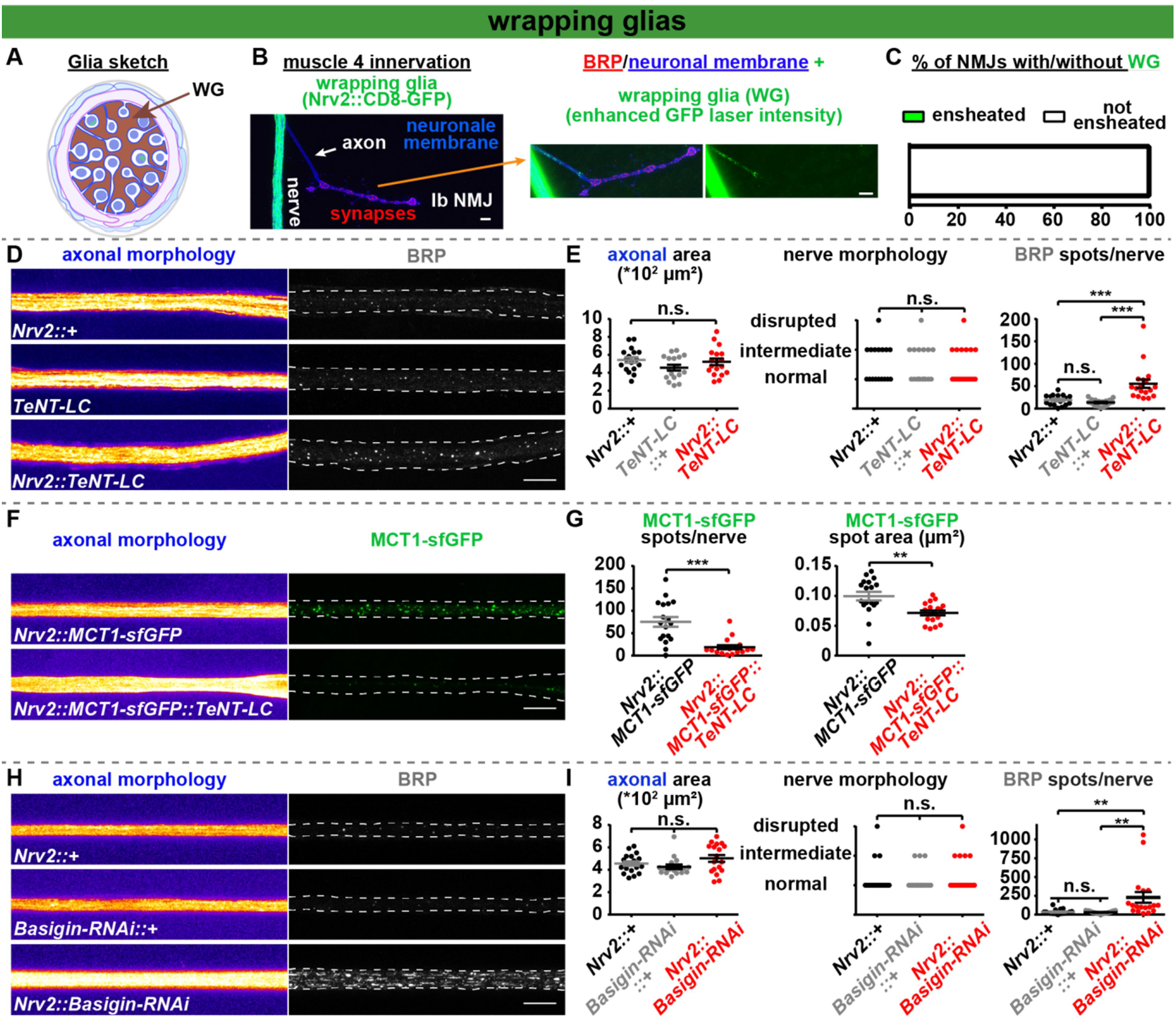
TeNT-LC action in WGs reduces nerve MCT1s numbers and causes axonal BRP accumulation similarly to Basigin-knockdown. **A** Cross section sketch of glial ensheathed motoneuron axon highlighting WGs (brown). **B,C** Example of muscle 4 NMJs of segments A2–A4 from 3^rd^-instar larvae expressing CD8-GFP in WGs (left) with enhanced GFP laser intensity (right). **C** Quantification of % of NMJs with/without WG ensheatment. **D,E** Nerves of segments A2–A4 (D) and quantification of axonal area, nerve morphology and BRP spots per nerve (E) from 3^rd^-instar larvae of *Nrv2::*+ (black), *TeNT-LC::*+ (grey) and *Nrv2::TeNT-LC* (red) animals. **F,G** Nerves of segments A2–A4 (F) and quantification of MCT1-sfGFP spots per nerve and spot area (G) from 3^rd^-instar larvae of *Nrv2::MCT1-sfGFP* (black) and *Nrv2::MCT1-sfGFP::TeNT-LC* (red) animals. **H,I** Same as in D,E but for *Nrv2::* + (black), *Basigin-RNAi::* + (grey) and *Nrv2::Basigin-RNAi* (red) animals. Dashed line in D, F, H indicates nerve area. Scale bars: (B) 5 μm; (D,F,H) 10 μm. Statistics: parametric one-way analysis of variance (ANOVA) test, followed by Tukey’s multiple comparison test except for (E; I: nerve morphology) where a non-parametric Kruskal-Wallis test and (G, MCT1-sf-GFP spots/nerve) where a Mann-Whitney U test and (G, MCT1-sfGFP spot area) where a Student’s t test was performed. ***p ≤ 0.001; n.s. (not significant) p > 0.05. (B,C) 23 NMJs, six larvae; (D, E) *Nrv2::*+: 17 nerves, six larvae; *TeNT::*+: 17 nerves, six larvae; *Nrv2::TeNT:* 18 nerves, six larvae; (F, G) *Nrv2::MCT1-sfGFP:* 18 nerves, six larvae; *Nrv2::MCT1-sfGFP::TeNT-LC:* 18 nerves, six larvae; (H,I) *Nrv2::* +: 17 nerves, six larvae; *Basigin-RNAi::*+: 17 nerves, five larvae; *Nrv2::Basigin-RNAi:* 18 nerves, six larvae. All panels show mean ± s.e.m. See also Fig. S3.

We hypothesized that impaired transport of synaptic BRP could be due to a shortage in metabolic energy in the neuron which is known to depend on glial cells (Edgar et al., 2009; Edgar et al., 2004; Griffiths et al., 1998; Nave, 2010). Here especially the Astrocyte-Neuron Lactate Shuttle (ANLS) Hypothesis (Pellerin and Magistretti, 1994) states that glial cells energetically support neurons by shuttling alanine and lactate via monocarboxylate transporters (MCTs) to fuel neuronal mitochondria in *Drosophila* and mice (Funfschilling et al., 2012; Lee et al., 2012; Liu et al., 2017; Machler et al., 2016; Pierre and Pellerin, 2005). We speculated that Syb was required to deliver MCTs to the plasma membrane and that TeNT-LC interfered with this. To test this hypothesis we co-expressed TeNT with an genomic GFP-tagged version of MCT1 (Sarov et al., 2016) which led to a severe reduction in the number and size of MCT1 positive spots along the peripheral nerve (Fig. 4F,G). The *Drosophila* genome harbors 15 putative MCT transporters (Gonzalez-Gutierrez et al., 2019). To further evaluate our hypothesis by an independent means we interfered with MCT function in general by investigating whether WG knockdown of Basigin, a mandatory accessory protein for the functional integration of multiple MCTs (Halestrap and Wilson, 2012), induced similar effects. Indeed and reminiscent of the effects observed upon TeNT-LC expression in WGs, WG specific expression of Basigin-RNAi led to a strong accumulation of axonal BRP (6-fold increase, Fig. 4H,I, slightly stronger than WG TeNT-LC: ~3-fold (Fig. 4D,E)) without any effect on axonal area, nerve integrity or neuronal lamella (Fig. 4H,I; Fig. S3C-E). The effect was specific to WG, as Basigin-knockdown in SPGs was without effect (Fig. S3F,G). Thus, our data are consistent with a crucial role of Syb in WGs to energetically supply axonal transport.

In conclusion we found unexpected and profound effects of TeNT-LC action on distinct glial cell types causing motoneuronal hyperexcitability, paralysis and death, and thus typical signs of TeNT-LC intoxication in mammals. Additionally we discovered disintegration of nerves, disruption of SJs, loss of nerve MCTs, glial accumulation of peptidergic vesicles and impaired axonal transport. We show that Syb, but not nSyb, is the relevant target and furthermore identify a differential requirement in two subpopulations of glial cells. While Syb appears to essentially contribute to SJ formation in SPG, it mediates neural metabolic support in WG. The observed phenotypes open new research avenues on how SNARE-mediated reactions in glial cells support neurons and other glial cells for proper nervous system function and whether disruptions of these contribute to diseases of peripheral motor control, including tetanus.

## Acknowledgements

We thank C. Klämbt, Stephan J. Sigrist and Sean Sweeney for comments on the manuscript and C. Klämbt for NrxIV-GFP and NrxIV-RNAi fly lines. We thank Katherine Cuthill, Björn von Domarus and Christina Fefler for help with initial experiments, Björn von Domarus for help with confocal analysis and Kiana Nafarieh for help with video analysis. We thank Andreas T. Grasskamp for his help with the orthogonal view. This work was supported by grants from the Deutsche Forschungsgemeinschaft to A.M. Walter (project ID 261020751 – Emmy Noether programme and Project-ID 278001972 – TRR 186).

## Author contributions

M.A.B and A.M.W conceived the project. M.A.B., K.P. and A.W.M. performed fly husbandry and maintenance. M.A.B. performed larvae behavior experiments and A.W.M. analyzed the data. M.A.B. and K.P. performed confocal experiments and analyzed the data. M.A.B., M.B. and A.W.M. performed electrophysiological experiments and M.A.B. and A.W.M. analyzed the data. M.A.B. and A.M.W wrote the paper with input from A.W.M..

## Methods

### Contact for Reagent and Resource Sharing

Further information and requests for resources and reagents should be directed to and will be fulfilled by the Lead Contact, Alexander M. Walter.

### Experimental Model and Subject Details

#### Fly husbandry, stocks and handling

Fly strains were reared under standard laboratory conditions (Sigrist et al., 2003) and raised at 25°C on semi-defined medium (Bloomington recipe). For RNAi experiments larvae were kept at 29°C. For experiments both male and female 3 instar larvae were used. The following genotypes were used: Figure 1 and Videos 1-3: Wild-type: +/+ (*w^1118^*). Repo::+: *Repo-Gal4*/+; TeNT-LC::+: *UAS-TeNT-LC*/+; Repo::TeNT-LC: *Repo-Gal4/UAS-TeNT-LC*. Figure 2: Repo::+: *Repo-Gal4*/+; nSyb-RNAi::+: *UAS-nSyb-RNAi*/+; Repo::nSyb-RNAi: *UAS-nSyb-RNAi*/+;*Repo-Gal4/*+; Syb-RNAi: *UAS-Syb-RNAi*/+; Repo::Syb-RNAi: *UAS-Syb-RNAi/+;Repo-Gal4/+*. Figure 3, Videos 4, 5 and 7: rL82::CD8-GFP: *rL82-Gal4/+; UAS-mCD8-GFP/+;* rL82::+: *rl82-Gal4/+;* rL82::TeNT-LC: *rL82-Gal4/+;UAS-TeNT-LC/+;* rL82::TeNT-LC:: Syb-GFVW: *rL82-Gal4/+; UAS-TeNT-LC/UAS-Syb-IsoA-QFVW;* rL82::NrxIV-GFP: *rl82-Gal4/+; NrxIV::GFP^454^/+;* rL82::NrxIV-GFP::TeNT-LC: *rl82-Gal4/+; NrxIV::GFP^454^/UAS-TeNT-LC;* NrxIV-RNAi::+: *UAS-NrxIV-RNAi/+;* rL82::NrxIV-RNAi: *rL82-Gal4/UAS-NrxIV-RNAi*. Video 6: rL82::TeNT-LC: *rL82-Gal4/(Wim); UAS-TeNT-LC/+;* Control: w1118. Figure 4: Nrv2::CD8-GFP: *Nrv2-Gal4/+; UAS-mCD8-GFP/+;* Nrv2::+: *Nrv2-Gal4/+;* TeNT-LC::+: *UAS-TeNT-LC/+;* Nrv2::TeNT-LC: *Nrv2-Gal4/+;UAS-TeNT-LC/+*; Nrv2:: MCT 1-sfGFP: *Nrv2-Gal4/+;MCT1-sfGFP/+*; Nrv2::MCT1-sfGFP::TeNT: *Nrv2-Gal4/+;MCT1-sfGFP/UAS-TeNT-LC;* Basigin-RNAi::+: *UAS-Basigin-RNAi/+;* Nrv2::Basigin-RNAi: *Nrv2-Gal4/+;UAS-Basigin-RNAi/+*. Fig. S1: rL82::+: *rl82-Gal4/+;* Syb-RNAi::+: *UAS-Syb-RNAi/+;* rL82::Syb-RNAi: *rL82-Gal4/UAS-Syb-RNAi;* TeNT-LC::+: *UAS-TeNT-LC/+;* rL82::TeNT-LC: *rL82-Gal4/+;UAS-TeNT-LC/+*. Fig. S2: rL82::CD8-GFP: *rL82-Gal4/+; UAS-mCD8-GFP/+;* rL82::CD8-GFP::TeNT-LC: *rL82-Gal4/+; UAS-mCD8-GFP/UAS-TeNT-LC;* rL82::CD8-GFP::Syb-RNAi: *rL82-Gal4/Syb-RNAi; UAS-mCD8-GFP/+;* rL82::+: *rl82-Gal4/+;* NrxIV-RNAi::+: *UAS-NrxIV-RNAi/+;* rL82::NrxIV-RNAi: *rL82-Gal4/UAS-NrxIV-RNAi;* rL82::ANF-EMD: *rL82-Gal4/+; UAS-ANF-EMD/+;* rL82::ANF-EMD::Syb-RNAi: *rL82-Gal4/Syb-RNAi; UAS-ANF-EMD/+*. Fig. S3: Nrv2:: CD8-GFP: *Nrv2-Gal4/+; UAS-mCD8-GFP/+*; Nrv2::CD8-GFP::TeNT-LC: *Nrv2-Gal4/+; UAS-mCD8-GFP/UAS-TeNT-LC;* Nrv2::+: *Nrv2-Gal4/+;* Basigin-RNAi::+: *UAS-Basigin-RNAi/+;* Nrv2::Basigin-RNAi: *Nrv2-Gal4/+;UAS-Basigin-RNAi/+;* rL82::+: *rL82-Gal4/+;* Basigin-RNAi::+: *UAS-Basigin-RNAi/+;* rL82::Basigin-RNAi: *rL82-Gal4/+;UAS-Basigin-RNA i/+*.

Stocks were obtained from: Repo-Gal4 (Sepp et al., 2001); UAS-TeNT-LC (Sweeney et al., 1995); UAS-nSyb-RNAi (VDRC #104531/#49201); UAS-Syb-RNAi (VDRC #102922); rL82-Gal4 (Sepp and Auld, 1999); Nrv2-Gal4 (Sun et al., 1999); NrxIV::GFP^454^ (Edenfeld et al., 2006); UAS-mCD8-GFP (Lee and Luo, 1999); MCT1-sf-GFP ((Sarov et al., 2016); VDRC #318191); UAS-Basigin-RNAi (VDRC #43307); UAS-NrxIV-RNAi (VDRC #8353); UAS-ANF-EMD (Rao et al., 2001).

## Method details

### Generation of UAS-Syb-IsoA-QFVW

UAS-Syb-IsoA-QFVW was generated by WellGenetics Inc. (Taipei, Taiwan). To generate cDNA encoding UAS-Syb-IsoA-QFVW, the sequence was amplified from cDNA clone SD05285 (obtained from DGRC). Point mutations were generated using the following primers:

Syn-RA-5’-QFVW:
F: gatctgcggccgcggctcgagATGGAGAACAACGAAGCCCC
R: GCTGCTCCCACACGGATGCTCCCTGCTCCAG
Syn-RA-3’-QFVW:
F: AGCATCCGTGTGGGAGCAGCAGGCCGGCAA
R: tcctctagaggtaccctcgagTTAGTGCGGCGGTGCTTG
PCR fragments were then cloned into into pUAST-attB vector using XhoI restriction sites. Generation of transgenic DNA micro-injection into embryos was performed by WellGenetics Inc., Taiwan using the PhiC31 integration system. The construct was inserted into strain 9725 (Bloomington, IN, USA): y[1] w[1118]; PBac{y[+]-attP-9A}VK00005.

### Immunostaining

Third-instar w1118 larvae were put on a dissection plate with both ends fixed by fine pins. Larvae were then covered by 50 μl of ice-cold hemolymph-like saline solution (HL3, pH adjusted to 7.2 (Stewart et al., 1994): 70 mM NaCl, 5 mM KCl, 20 mM MgCl_2_, 10 mM NaHCO_3_, 5 mM Trehalose, 115 mM D-Saccharose, 5 mM HEPES). Using dissection scissors a small cut at the dorsal, posterior midline of the larva was made from where on the larvae was cut completely open along the dorsal midline until its anterior end. Subsequently, the epidermis was pinned down and slightly stretched and the internal organs and tissues removed. Care was taken not to harm the ventral nerve cord and the peripheral nerves. The dissected samples were washed 3x with ice-cold HL3 and then fixed for 5 minutes with icecold methanol. After fixation, samples were briefly rinsed with HL3 and then blocked for 1h in 5% native goat serum (NGS; Sigma-Aldrich, MO, USA, S2007) diluted in phosphate buffered saline (Carl Roth, Germany) with 0.05% Triton-X100 (PBT). Subsequently dissected samples were incubated with primary antibodies (mouse Nc82 = anti-BRP^C-term^ (1:100, Developmental Studies Hybridoma Bank, University of Iowa, Iowa City, IA, USA; AB Registry ID: AB_2314865); rabbit BRP^Last200^ (1:1000; (Ullrich et al., 2015)); rabbit GluRIID (1:500; (Qin et al., 2005)); mouse GFP 3E6 (1:500, Thermo Fisher Scientific Inc., MA, USA, A-11120; AB Registry ID: AB_221568)) diluted in 5% NGS in PBT overnight. Afterwards samples were washed 5x for 30 min with PBT and then incubated for 4h with fluorescence-labeled secondary antibodies (goat anti-HRP-647 (1:500, Jackson ImmunoResearch 123-605-021, PA, USA); goat anti-rabbit-Cy3 (1:500, Jackson ImmunoResearch 111-165-144, PA, USA); goat anti-mouse-Cy3 (1:500, Jackson ImmunoResearch 115-165-146, PA, USA); anti-Phalloidin-Atto565 (1:1700; Sigma-Aldrich, MO, USA, 94072); goat anti-mouse Alexa-Fluor-488 (1:500, Life Technologies A11001, CA, USA)) diluted in 5% NGS in PBT. Samples were then washed overnight in PBT and subsequently mounted in vectashield (Vector labs, CA, USA) on microscope slides (Carl Roth, Germany; H868) and sealed with coverslips (Carl Roth, Germany, H 875). Antibodies obtained from the Developmental Studies Hybridoma Bank were created by the NICHD of the NIH and maintained at The University of Iowa, Department of Biology, Iowa City, IA 52242.

### Image Acquisition, Processing, and Analysis

Confocal microscopy was performed with a Leica SP8 microscope (Leica Microsystems, Germany). Images were acquired at room temperature. Confocal imaging was done using a 63×1.4 NA oil immersion objective with a zoom of 1.8 and z-step size of 0.25 μm. All confocal images were acquired using the LAS X software (Leica Microsystems, Germany). Images from fixed samples were taken from nerve bundles exiting the ventral nerve cord of segments A2–A4 (see Figure 1A for depicted ROI). Confocal stacks were processed with ImageJ software (http://rsbweb.nih.gov/ij/). Quantifications of axonal BRP spot numbers were performed following an adjusted manual (Andlauer and Sigrist, 2012), briefly as follows. The signal of a HRP-647 antibody was used as template for a mask, restricting the quantified area to the shape of the nerve. The original confocal stacks were converted to maximal projections and a mask of the axonal BRP spots was created by applying a threshold to remove irrelevant lower intensity pixels. The threshold was adjusted manually and individually to every image to detect all axonal BRP spots. The segmentation of single spots was done semi-automatically via the command “Find Maxima” embedded in the ImageJ software and by hand with the pencil tool and a line thickness of 1 pixel. To remove high frequency noise a Gaussian blur filter (0.5 pixel Sigma radius) was applied. The processed picture was then transformed into a binary mask using the same lower threshold value as in the first step. This binary mask was then projected onto the original unmodified image using the “min” operation from the ImageJ image calculator. BRP spot numbers and sizes were determined using the “Analyze particles” function (particle size > 2 pixels) embedded into ImageJ. Line profiles were measured using the plot profile function of ImageJ. To measure the axonal (HRP) or lamellar (GluRIID) area, the signal of the HRP-647 or the GluRIID antibody was used as template for a binary mask. The area was then quantified using the “wand tool” to select the area and “measure” function embedded into ImageJ to quantify the area. For orthogonal sections, confocal RGB stacks were processed using MATLAB R2016a as follows: first, the whole stack containing the three channels (BRP, GluRIID, HRP) was loaded using the command “imread” in a loop iterating through the first to last image of the stack. Then, the three channels were separated into three stacks for the following permutation operation. Using the command “permute”, the 2^nd^ and 3^rd^ dimension were switched separately in each stack. Lastly, each of the three permuted stacks was written to a file using the command “imwrite” in a loop iterating through all images of the stack, resulting in three .tif-files (each for one channel) showing the orthogonal view of the imaged nerve. The respective code is available upon request. Orthogonal views of each nerve (BRP, GluRIID, HRP) were then merged using ImageJ.

To evaluated nerve disruption, nerves were manually categorized by their degree of disintegration. Category 1: normal morphology: long, thin and smooth nerves (see control images in Fig. 1E for example); category 2: intermediate: slight morphological alterations like single defasciculating axons, slightly increased nerve area; category 3: disrupted: complete disintegration of the nerve integrity, defasciculation of whole parts of the nerve, large expansion of the nerve area (see *Repo::Syb-RNAi* image in Fig. 2E for example).

Images for figures were processed with ImageJ software to enhance brightness using the brightness/contrast function.

### Video of larval and adult fly movement

Larval behavior was investigated by placing 3^rd^ instar larvae of the correct genotype in the center of a petri dish (diameter: 5.4 cm) that contained two ml of water to avoid larvae attachment to the plastic dish or larvae crawling up the wall of the dish. Videos were recorded with a Samsung Galaxy A50 for ca. 90s using the imbedded video function. A ruler was placed next to the dish to allow measuring the walking distance. For analysis, videos were converted to AVI format and compressed using FFmpegTool software (v4.3, ffmpeg.zeranoe.com). Videos were analyzed by a different person than the one recording the video blinded for genotype. Using Fiji software (ImageJ 1.51n) a substack was created for each video, selecting every 50^th^ frame. This reduced the framerate from 30 Hz to 0.6 Hz. Using the manual tracking plugin of Fiji, the head of the larva was selected and tracked in each frame of the substack for 80 s (Fig. 3D,E; Videos 4,5,7) or 85 s (Fig. 1C,D; Videos 1-3) after it was added to the dish. Final larval movement tracks were saved as ROIs and the total distance travelled by each larva calculated.

For adult behavior assessment (Video 6) two *rL82::TeNT-LC* flies and one control (w1118) were put in an empty food vial and illuminated with a ZLED CLS 600 (Zett Optics, Gemany) light source. To induce fly movement, the vial was occasionally banged on the table.

### Protein sequence alignment

Sequences of Syb-PA (ID: FBpp0087450) and nSyb-PA (ID: FBpp0072697) were obtained from flybase (flybase.org, version FB2020_02; date: May 28 2020). Alignment was performed using Clustal Omega ((Sievers et al., 2011); https://www.ebi.ac.uk/Tools/msa/clustalo/; version 1.2.4). Gonnet PAM250 matrix was used to compare sequence substitutions: * identical aa,: score >0.5,. score <0.5, “gap” score below 0.

### Electrophysiology

Third instar larvae were individually placed on Sylgard (184, Dow Corning, Midland, MI, USA) and pinned at the head and the tail. 40 μl modified hemolymph-like solution (HL3; (Stewart et al., 1994) composition (in mM): NaCl 70, KCl 5, MgCl_2_ 10, NaHCO_3_ 10, trehalose 5, sucrose 115, HEPES 5, CaCl_2_ 0, pH adjusted to 7.2) was pipetted onto the larva at room temperature (~22°C). A small incision was made with a sharp scissors in the dorsal cuticle near the tail pin. Starting from this posterior incision, a cut was made along the length of the larva extending beyond the head pin. The cuticle was pinned down twice on either side. The intestines and trachea were cut at the posterior and held firmly with a forceps as the remaining connections to the body were cut before being fully removed, taking care not to pull on the preparation. The brain was held slightly raised above the preparation and the segmental nerves cut without touching the underlying muscle, before finally removing the brain. The Sylgard block and completed larval preparation was placed in the recording chamber which was filled with 2 ml HL3 (plus 0.4 mM CaCl_2_, 10 mM MgCl_2_). Recordings were performed at room temperature in current clamp mode at muscle 6 in segments A2/A3 as previously described (Zhang and Stewart, 2010) using an Axon Digidata 1550A digitizer, Axoclamp 900A amplifier with HS-9A x0.1 headstage (Molecular Devices, CA, USA) and on a BX51WI Olympus microscope with a 40X LUMPlanFL/IR water immersion objective. Sharp intracellular recording electrodes were made using a Flaming Brown Model P-97 micropipette puller (Sutter Instrument, CA, USA) with a resistance of 20-35 MΩ and backfilled with 3 M KCl. Cells were only considered with a membrane potential of less than - 60 mV and membrane resistances greater than 3 MΩ. All recordings were acquired using Clampex software (v10.5), filtered with a 5 kHz low-pass filter and sampled at 10-50 kHz. eEPSPs were recorded by stimulating the appropriate nerve at 10 s intervals five times (8 V, 300 μs pulse) using an ISO-STIM 01D stimulator (NPI Electronic, Germany). Stimulating suction electrodes were pulled on a DMZ-Universal Puller (Zeitz-Instruments GmbH, Germany) and fire polished using a CPM-2 microforge (ALA Scientific, NY, USA). A maximum of two cells were recorded per animal.

Analysis was performed with Clampfit 10.5 and Graphpad Prism 6 software. mEPSPs were further filtered with a 500 Hz Gaussian low-pass filter. A single mEPSP template was generated for each 1-minute cell recording and used to identify individual mEPSPs, calculating the mean mEPSP amplitude per cell. To calculate eEPSP amplitudes, an average trace was generated from the five traces and the amplitude of the first response peak in the resulting average trace calculated. To determine hyperactivity or aberrant cell responses, a threshold was set at 8 mV. The number of response peaks above the 8 mV threshold for each cell was determined and normalized to the number of stimuli (five). Aberrant activity in a cell was defined as multiple response peaks above the 8 mV threshold following each individual stimulus, or any response to an individual stimulus below the 8 mV threshold including failures.

### Quantification and Statistical Analysis

Data were analyzed using Prism (GraphPad Software, CA, USA). Per default two-sided Student’s t test was performed to compare the means of two groups unless the data were non-normally distributed (as assessed D’Agostino-Pearson omnibus normality test) in which case they were compared by a two-tailed Mann-Whitney U test. For comparison of more than two groups, one-way analysis of variance (ANOVA) tests were used, followed by a Tukey’s multiple comparison test. In the case of ordinal data (nerve morphology) a Kruskal-Wallis test was used. Means are annotated ± s.e.m.. Asterisks are used to denote significance: *, p < 0.05; **, p < 0.01; ***, p < 0.001; n.s. (not significant), p > 0.05.

### Data and Software Availability

The data that support the findings of this study, additional information and requests for resources and reagents as well as Matlab or ImageJ codes used in this study are available from Alexander M. Walter upon request.

## Video titles and legends

**Videos 1-3: Pan-glial TeNT-LC expression severely impairs larval movement.** Video of larval movement over 85s for *Repo::*+ (Video 1), *TeNT::*+ (Video 2) and *Repo::TeNT* (Video 3). See also Figure 1C,D for analysis.

**Videos 4, 5 and 7: TeNT-insensitive Syb-variant recovers larval movement in SPG TeNT-LC expressing animals.** Video of larval movement over 80s for *rL82::*+ (Video 4), *rL82::TeNT* (Video 5) and *rl82::TeNT::Syb-QFVW* (Video 7). See also Fig. 3D,E for analysis.

**Video 6: TeNT-LC expression in SPGs severely paralyzes adult flies.** Video of two *rL82::TeNT-LC-LC* flies and one control. *rL82::TeNT-LC-LC* flies are severely paralyzed (uncoordinated or no movement; inability to turn around when laying on their backs; no reaction to banging) while the control fly rapidly moves upwards after bang-shock.

