## Supplementary Material for "Tetanus neurotoxin sensitive SNARE-mediated glial signaling limits motoneuronal excitability"

### Supplementary figure 1:

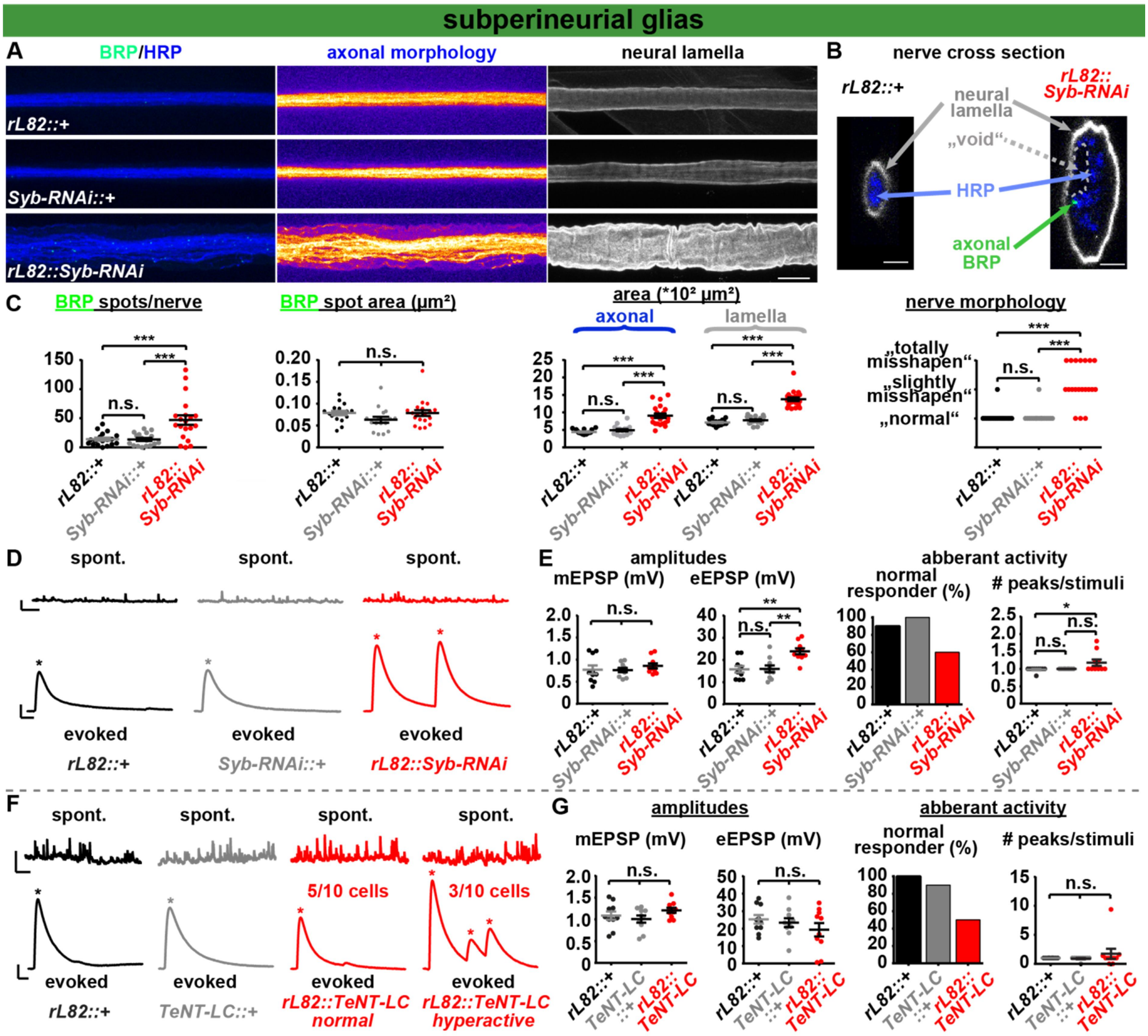


**Supplementary Fig. 1. Syb is targeted by TeNT-LC in SPGs. A-C** Nerves of segments A2–A4 (A; neural lamella visualized via GluRIID staining), orthogonal nerve cross section (B) and quantification of BRP spots per nerve, axonal BRP spot area, axonal/lamella area and nerve morphology (C) from 3rd-instar larvae of *rL82::+* (black), *Syb-RNAi::+* (grey) and *rL82:: Syb-RNAi* (red) animals. **D,E** Representative mEPSP (spont.) and eEPSP (evoked) traces (D) and quantification of mEPSP/eEPSP amplitudes, % of recorded cells displaying normal activity and the normalized number of response peaks per stimulus (E) of *rL82::+* (black), *Syb-RNAi::+* (grey) and *rL82:: Syb-RNAi* (red) animals. Asterisks (D) indicate postsynaptic response peaks (eEPSP). **F,G** Representative mEPSP (spont.) and eEPSP (evoked) traces (F; for *rl82::TeNT-LC* one normal and one hyperactive trace is shown) and quantification of mEPSP/eEPSP amplitudes, % of recorded cells displaying normal activity (single eEPSP in response to single stimulation) and the normalized number of response peaks per stimulus in *rL82::+* (black), *TeNT-LC::+* (grey) and *rL82::TeNT-LC* (red) animals (G). Asterisks indicate postsynaptic response peaks (eEPSP). Scale bars: (A) 10 µm; (B) 2.5µm; (D,F) mEPSP: 1 s, 2 mV;

eEPSP: 25 ms, 5 mV. Statistics: parametric one-way analysis of variance (ANOVA) test, followed by Tukey's multiple comparison test except for (C, nerve morphology) where a non-parametric Kruskal-Wallis test was performed. ***$p \leq 0.001$; **$p \leq 0.01$; *$p \leq 0.05$; n.s. (not significant) $p > 0.05$. (A-C) *rl82::+*: 18 nerves, six larvae; *Syb-RNAi::+*: 17 nerves, six larvae; *rL82::Syb-RNAi*: 20 nerves, six larvae; (D,E) *rl82::+*: 10 cells, five larvae; *Syb-RNAi::+*: 10 cells, five larvae; *rL82::Syb-RNAi*: 10 cells, five larvae; (F,G) *rL82::+:* 10 cells, five larvae; *TeNT::+:* 10 cells, five larvae; *rL82::TeNT:* 10 cells, five larvae. All panels show mean ± s.e.m.. See also Fig. 3.

**Supplementary figure 2:**

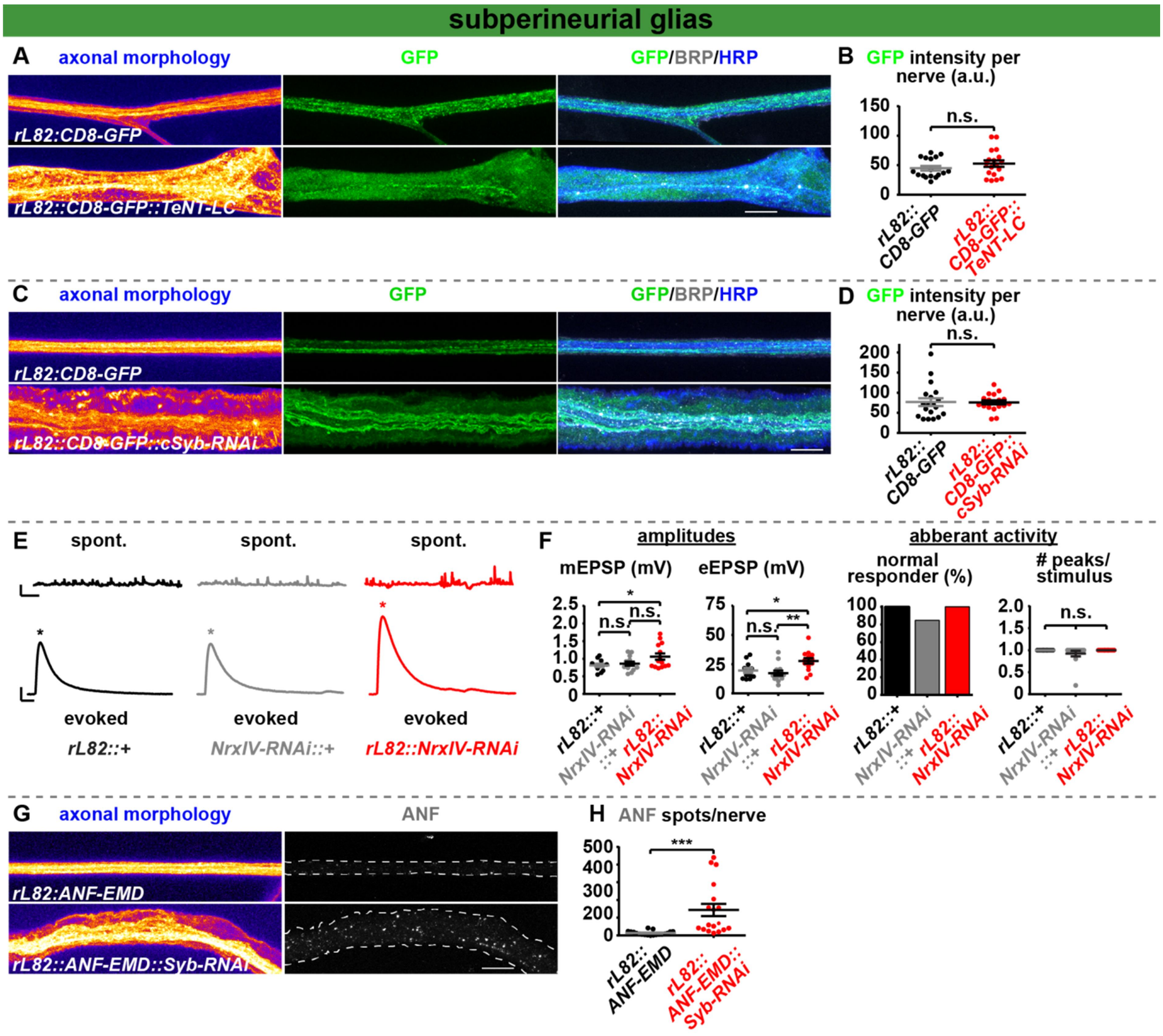


**Supplementary Fig. 2: TeNT-LC/Syb-RNAi expression in SPGs does not kill SPG cells but causes ANF accumulation. A,B** Nerves of segments A2–A4 (A) and quantification of GFP intensity per nerve area (B) from 3rd-instar larvae of *rL82::CD8-GFP* (black) and *rL82::CD8-GFP::TeNT-LC* (red) animals. **C,D** same as in A,B but for *rL82:: CD8-GFP* (black) and *rL82::CD8-GFP::Syb-RNAi* (red) animals. **E,F** Representative mEPSP (spont.) and eEPSP (evoked) traces (E) and quantification of mEPSP/eEPSP amplitudes, % of recorded cells displaying normal activity (single eEPSP in response to single stimulation) and the normalized number of response peaks per stimulus (F) for *rL82::+* (black), *NrxIV-RNAi::+* (grey) and *rL82::NrxIV-RNAi* (red) animals. Asterisks indicate postsynaptic response peaks (eEPSP). **G,H** Nerves of segments A2–A4 (G) and quantification of ANF spots per nerve (H) from 3rd-instar larvae of *rL82::ANF-EMD* (black) and *rL82::ANF-EMD::TeNT* (red) animals. Scale bars: (A,C,G) 10 µm; (E) mEPSP: 1 s, 2 mV; eEPSP: 25 ms, 5 mV. Statistics: Student's t test (B,H), Mann-Whitney U test (D) and parametric one-way analysis of variance (ANOVA) test, followed by Tukey's multiple comparison test (F). ***$p \leq 0.001$; **$p \leq 0.01$; *$p \leq 0.05$; n.s. (not significant) $p > 0.05$. (A,B) *rl82::CD8-GFP*: 17 nerves, six larvae; *rl82::CD8-*

*GFP::TeNT*: 18 nerves, six larvae; (C,D) *rl82::CD8-GFP*: 19 nerves, six larvae; *rl82::CD8-GFP::Syb-RNAi*: 18 nerves, six larvae. (E,F) *rL82::+:* 12 cells, seven larvae; *NrxIV-RNAi::+:* 13 cells, seven larvae; *rL82::NrxIV-GFP:* 14 cells, seven larvae; (G,H) *rL82::ANF-EMD*: 19 nerves, six larvae; *rL82::ANF-EMD::Syb-RNAi*: 18 nerves, six larvae. All panels show mean ± s.e.m.. See also Fig. 3.

**Supplementary figure 3:**

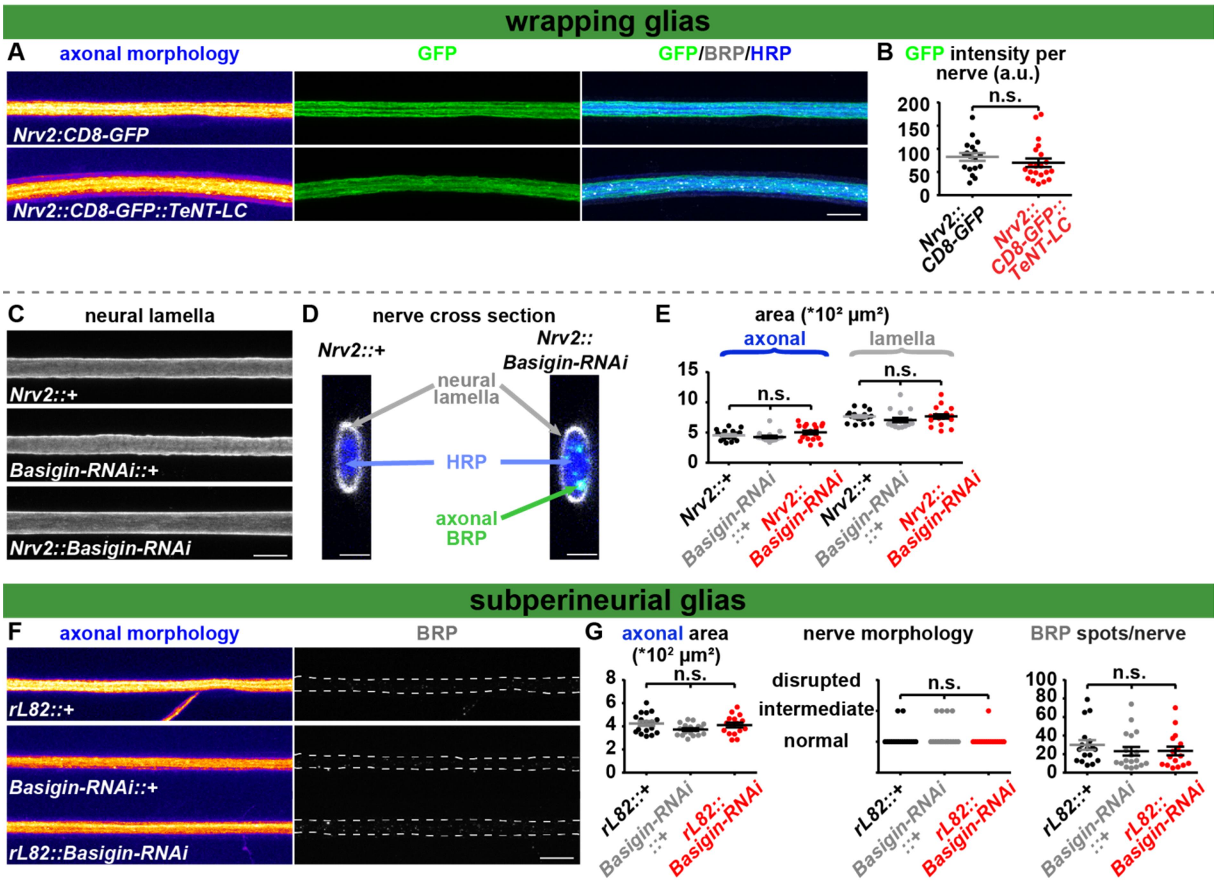


**Supplementary Fig. 3. Neither TeNT-LC nor Basigin-RNAi expression in WG affect nerve morphology. A,B** Nerves of segments A2–A4 (A) and quantification of GFP intensity per nerve area (B) from $3^{rd}$-instar larvae of *Nrv2::CD8-GFP* (black) and *Nrv2::CD8-GFP::TeNT-LC* (red) animals. **C-E** Nerves of segments A2–A4 (C; neural lamella visualized via GluRIID staining), orthogonal nerve cross section (D) and quantification of axonal/lamella area (E) from $3^{rd}$-instar larvae of *Nrv2::+* (black), *Basigin-RNAi::+* (grey) and *Nrv2::Basigin-RNAi* (red) animals. **F-G** Nerves of segments A2–A4 (F) and quantification of axonal area, nerve morphology and BRP spots per nerve (G) from $3^{rd}$-instar larvae of *rL82::+* (black), *Basigin-RNAi::+* (grey) and *rL82::Basigin-RNAi* (red) animals. Scale bars: (A, C, F) 10 µm; (D) 2.5 µm. Statistics: parametric one-way analysis of variance (ANOVA) test, followed by Tukey's multiple comparison test except for (B) where a Mann-Whitney U Test and (G, nerve morphology) where a non-parametric Kruskal-Wallis test was performed. ***$p \leq 0.001$; **$p \leq 0.01$; *$p \leq 0.05$; n.s. (not significant) $p > 0.05$. (A,B) *Nrv2::CD8-GFP*: 18 nerves, six larvae; *Nrv2::CD8-GFP::TeNT*: 21 nerves, six larvae; (C-E) *Nrv2::+*: 17 nerves, six larvae; *Basigin-RNAi::+:* 17 nerves, five larvae; *Nrv2::Basigin-RNAi*: 18 nerves, six larvae; (F,G) *rL82::+*: 17 nerves, six larvae; *Basigin-RNAi::+:* 18 nerves, six larvae; *rL82::Basigin-RNAi*: 18 nerves, six larvae. All panels show mean ± s.e.m.. See also Fig. 4.
